# Intratumoral expression of IL-12 and CD40 ligand (CD154) from plasmids generates antitumor responses that eliminate tumoral T regs

**DOI:** 10.1101/2025.05.20.655082

**Authors:** Gregory W. Ho, Alicia Santos, Jennifer Fields, Pamela C. Rosato, Mary Jo Turk, Nicole F. Steinmetz, Steven Fiering

## Abstract

Intratumoral immunotherapy (ITIT) strives to generate effective antitumor immunity by directly stimulating the immune system in tumors to reverse local tumor-mediated immune suppression. *In vivo* expression of Interleukin-12 (IL-12) using *in vivo* plasmid transfection as an intratumoral cancer immunotherapy entered Phase II clinical trials for metastatic melanoma but to limited clinical success. We sought to improve the efficacy of *in vivo* IL-12 electroporation by the addition of a CD154 (CD40 ligand)- expressing plasmid to the IL-12 encoding plasmid treatment and assessing efficacy against solid tumors. Mice with intradermal B16F10 melanoma or MC38 murine colon carcinoma tumors received 2 weekly intratumoral (IT) injections of plasmids encoding IL-12 and CD154, followed by *in vivo* electroporation. The addition of CD154 to IL-12 was superior to IL-12 alone and resulted in frequent tumor clearance of treated tumors, marked by an increase in CD8 T cells and a drastic reduction in T regulatory cells in the tumor microenvironment. Tumor treatment responses were abrogated in mice which lack conventional DC1 cells (BatF3 KO) or lack CD8 T cells. These findings highlight the potential of adding CD154 to IL-12 plasmid electroporation as a cancer immunotherapy and suggest that other combinations would be therapeutically valuable.

## Introduction

Any effective anti-tumor immunotherapy must overcome the immune suppressive tumor microenvironment (TME).[1,2] Robust anti-tumor immunity relies on an activated and functional CD8 T cell compartment, and recent advances in cancer immunotherapies aim to generate effective T cell responses against tumor cells. [3] There are many new cancer immunotherapies under investigation, including intratumoral immunotherapies (ITIT). ITIT delivers immune stimulating treatments or reagents directly into one or more tumors to shift the suppressive tumor microenvironment (TME) into a proinflammatory TME and stimulate an effector T cell response within the tumor against the whole repertoire of tumor antigens. [4] ITIT uses high local but low systemic doses of reagents, making it generally safer and less expensive than systemic immunotherapy. Additionally, it can be accomplished rapidly with standard, non-personalized reagents and does not need tumor antigen identification because any antigens expressed by the tumor are potential targets and are selected by the immune system. While not yet being used as neoadjuvant treatment in the clinic, the treatment is generally rapid and the immune impacts are established quickly, so it could be used when metastatic disease is suspected but not identified. There are recently approved ITIT approaches, including oncolytic viruses (TVEC) and Rose Bengal (PV-10), however ITIT development is still minimal. Fundamentally, radiation therapy is a weak ITIT that led to the identification of “abscopal effects”. [4–6]

We have utilized a plant virus, cowpea mosaic virus, a multi-toll-like receptor (TLR) agonist, as an immune adjuvant in ITIT that offers robust protection in mouse models and companion dog cancer patients of both the treated tumor and distal non-treated tumors, (abscopal effects). CPMV treatment relies on various combinations of neutrophils, APCs, adaptive immune cells, IL-12, type I interferons and interferon (IFN)-y. [7–9] The addition of agonistic CD40 antibody along with CPMV into the tumor increased cDC1 activation and priming of CD8 T cell responses, inducing a greater anti-tumor response both locally and systemically, and focusing our interest on CD40 agonism for ITIT. [9]

*In vivo* plasmid electroporation is a promising platform for ITIT. [10] Expression plasmids are injected into the tumor and subsequently electroporated, inducing the uptake and expression of the encoded immune stimulatory molecules. [10] The best studied cytokine in plasmid electroporation is Interleukin 12 (IL-12), a proinflammatory cytokine, which stimulates T cells to an effector fate and promotes effector function and interferon gamma production. [11,12] IL-12 electroporation offers a targeted and prolonged expression of IL-12, which allows an increase in local T cell number and effector function, while circumventing the potential toxicities associated with systemic delivery of cytokines.[13] IL-12 electroporation studies have observed elevated CXCR3 gene signature in CD8 T cells in treated tumors, corresponding with the increase in CD8 T cell infiltration.[14] In addition to increased overall tumoral CD8 T cells, IL-12 increases the frequency of tumor antigen-specific CD8 T cells contributing to increased local and systemic antitumor immunity. [15] Furthermore, the requirement of CD8 T cells is highlighted by impact of CD8 T cell depleting antibody, which nearly abolishes efficacy of IL-12 electroporation.[16]

*In vivo* electroporation is feasible in humans and there is an FDA approved electroporation device [17], although it is generally used for irreversible electroporation to directly kill cells in the TME rather than to transfect cells *in vivo*. *Tavokinogene* is an IL-12 electroporation therapeutic that has gone into Phase II clinical trials for metastatic melanoma. While it has shown minimal clinical benefit, it highlights the feasibility for plasmid electroporation as a cancer therapeutic. [18–20] Subsequent plasmid electroporation studies have utilized IL-12 as a platform in combination with other cytokines, aimed at delivering additional T cell stimulus to the TME, including IL-2, IL-15, membrane-anchored CD3, and CXCL9. [16,21–24] While most of these molecules offer some reduction in tumor burden as a monotherapy, their combination with IL-12 has better efficacy than either protein alone. [11] With this in mind, we sought to potentiate IL-12s immune impact by expressing IL-12 and causing CD40 agonism through IL-12 and CD154 (CD40 ligand) *in vivo* plasmid electroporation.

Previous studies showed that activation of CD40 provides potent maturation and anti-apoptotic signals to DCs and other antigen-presenting cells (APC), leading to the upregulation of major histocompatibility complex (MHC) molecules and increased costimulatory markers such as CD86. [25] A CD40-activated DC population amplifies and expands the CD8 T cell compartment through efficient priming and secretion of stimulatory cytokines such as IL-12, polarizing the adaptive immune response toward a Th1 response. Critically, CD40 stimulation appears uniquely capable of inducing the expression of TNF superfamily members on DCs, including CD70, CD134 (OX40 ligand) and CD137 (4-1BB ligand) which provide critical signaling that supports the continued expansion of effector T cells and priming an anti-tumor response. [26,27] Clinically, systemic administration of agonistic CD40 antibody as a monotherapy generates minimal tumor response. The one exception is selicrelumab, that produced objective partial responses in 27% of patients with advanced melanoma in a first-in-human single-dose study, although nonmelanoma solid tumors did not respond. [28] However, significant adverse effects with systemic transfusions have also been observed, dissuading development of agonistic CD40 monoclonal antibody as an immunotherapy. [29] Therefore, in order the avoid the systemic toxicities associated with systemic delivery of CD40 agonism, we investigated the addition of a CD154- expressing plasmid to enhance IL-12 plasmid electroporation ITIT.

## Results

### The addition of CD154 to IL-12 electroporation eliminates established treated tumors

To determine the effect of adding CD154-expressing plasmid to the IL-12 expressing plasmid electroporation platform, mice bearing intradermal B16F10 melanoma tumors were injected with non-integrating plasmids expressing either an irrelevant plasmid (expressing β-Geo, a fusion protein of E. coli beta-galactosidase and neomycin), CD154 expressing plasmid, IL-12-expressing plasmid or both. ITIT electroporation treatments of established tumors (50mm^3^) were applied weekly for 2 weeks. While CD154 and IL-12 alone each reduced the growth rate of treated B16F10 tumors, none of the tumors were eliminated and there was minimal improved survival. However, the combination eradicated 80% of intradermal B16F10 flank tumors with no recurrence observed (monitored until Day 40) (**Figure 1A**). Similarly, we evaluated the efficacy of the combination in another tumor model, MC38 colon carcinoma and observed similar results in MC38 tumors (**Figure 1B**), demonstrating that CD154 + IL-12 efficacy is not limited to B16F10. Interestingly, in a B16F10 two-tumor model, with one treated tumor and an untreated tumor on the opposite flank to model abscopal effects, the additive effect of the CD154 to IL-12 electroporation was limited to the treated tumor only. Both IL-12 and CD154 + IL-12 reduced growth of untreated tumors but tumor growth did not differ between the two treatment groups, indicating CD154 is not increasing IL-12 driven systemic protection (**Supplementary Figure 1A, B)**.

**Figure 1.**
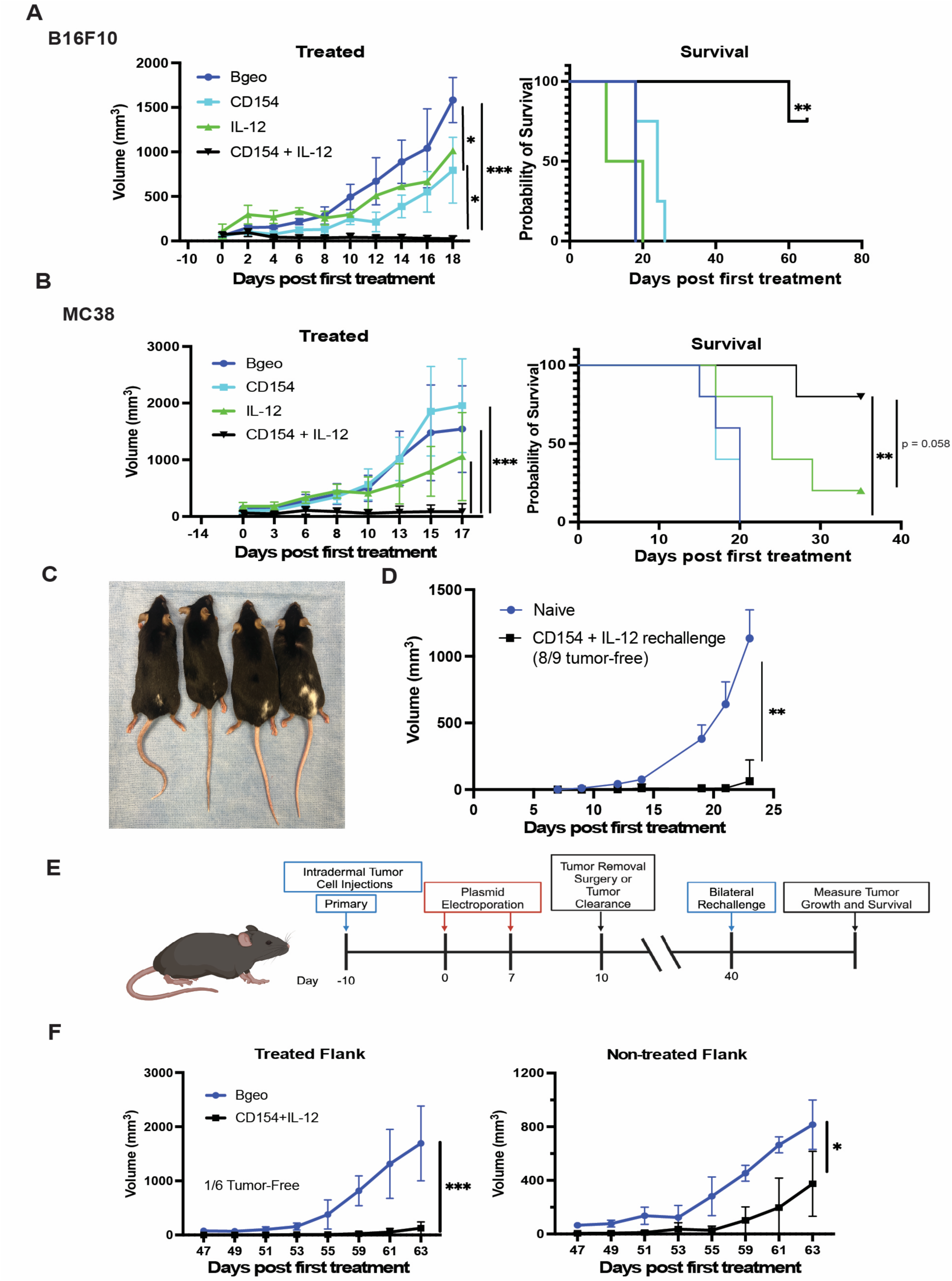
CD154 and IL-12 electroporation improves anti-tumor efficacy on the treated tumor. A) B16F10-bearing mice treated twice with CD154 + IL-12 electroporation abolish most treated intradermal tumors. B) MC38 colon carcinoma-bearing mice exhibit similar high frequency clearance of intradermal tumors. C) Mice that cleared B16F10 melanoma following treatment with CD154 + IL-12 electroporation exhibit localized vitiligo at the tumor site. D) CD154 + IL-12 treated mice that cleared tumors rejected tumors upon rechallenge on previously treated flank. E) Treatment timeline of B16F10 bilateral rechallenge experiment on CD154 + IL-12-treated mice that had cleared tumors or control plasmid-treated that had tumors surgically resected on Day 10. F) CD154 + IL-12 treated mice that had previously cleared B16F10 flank tumors, exhibited enhanced systemic protection against rechallenge as compared to mice that had B26F10 surgically removed on day 10. All experiments were repeated at least once with similar results. N=4-5. Growth curves were analyzed using two-way ANOVA. Survival curves were compared using log-rank (Mantel-Cox) test, with p > 0.05 as ns, p < 0.05 as *, p < 0.01 as **, and p < 0.001 as ***.

Mice that cleared B16F10 tumors with CD154 + IL-12 treatment exhibited local vitiligo, an autoimmune disorder that involves depigmentation of the fur by CD8 T cells, suggesting the emergence of a melanocyte-specific T cell populations (**Figure 1C**). [30] In order test immune memory, we rechallenged CD154+ IL-12-treated mice that had previously cleared treated tumors with B16F10 on the treated flank. At Day 40 post first treatment, we observed superior protection against rechallenge on the treated flank of the CD154 + Il-12 mice compared to naïve controls (**Figure 1D**). To compare CD154+ IL-12 memory against a B16F10 antigen-experienced mouse, we treated mice with either β-geo as a control or CD154 + IL-12. Mice that received β-geo had their tumors surgically removed at Day 10 (three days post second treatment), while the CD154 + IL-12 were allowed to clear tumors. On Day 40, mice were rechallenged bilaterally to determine CD154 + IL-12 systemic protection. CD154 + IL-12 treated mice displayed systemic protection on the non-treated as well as treated flanks (**Figure 1F**). This data highlights the potential of adding CD154 to IL-12 electroporation for improved anti-tumor efficacy and increased immune memory.

### Injection of irrelevant plasmid reduces tumor burden through TLR9 agonism

In the past, irreversible electroporation (IRE) has been evaluated as a possible cancer therapy. IRE is the application of short pulses of high voltage, up to 3kV, to induce cell death. [31] While our treatment parameters are less toxic to the tumor than IRE, we sought to determine the effect of electroporation alone on tumor growth. The application of electrical charge did not affect tumor growth in B16F10, however the administration of β-geo plasmid without electrical charge reduced tumor growth and seems to further impede growth with the delivery of charge. (**Supplementary Figure 2A**) To determine the mechanism behind reduced tumor growth in β-geo plasmid administration alone, we investigated the potential for passive plasmid uptake and expression utilizing the IL-12 plasmid and found that the application of electrical charge was required for increased expression, indicating the control plasmid itself was inducing a response without detectable protein expression. (**Supplementary Figure 2B).** It is well-established that many cells, including DCs express multiple TLRs including TLR9, which recognizes unmethylated CpG-DNA. TLR agonists can activate DCs, inducing upregulation of costimulatory molecules as well as inflammatory cytokines. [32] We investigated the effect of TLR9 agonism and found that the effect on tumor growth of plasmid injection without electroporation was eliminated in TLR9 knockout mice (**Supplementary Figure 2C, D**). Thus, indicating that the injection of plasmids (that lack CpG methylation), triggers TLR9 agonism in the TME, adding to the efficacy of the plasmid payloads.

### CD8 T cells drive CD154 + IL-12-induced anti-tumor response

To determine immune mechanisms behind CD154 + IL-12 electroporation, we utilized antibody depletions to identify immune cells required for efficacy. We observed that depletion of natural killer cells had a negligible effect on CD154 + IL-12 efficacy, however the depletion of CD8 T cells completely abrogated the efficacy of CD154 + IL-12 (**Figure 2A).** Similarly, we evaluated the mechanism behind IL-12 electroporation alone and found that it was also CD8 T cell-driven (**Figure 2B**). We observed an increase in IL-12 expression in treated tumors with IL-12 or CD154 + IL-12 electroporated tumors that corresponded with a concurrent increase in IFN-y in both groups (**Supplementary Figure 3A**). Following up, we utilized an IFN-y-neutralizing antibody to evaluate the effect on CD154 + IL-12 efficacy. Antibody neutralization of IFN-y erased CD154 + IL-12 anti-tumor effect (**Figure 2C**). All of this taken together, demonstrates that CD154 + IL-12 electroporation requires CD8 T cells and subsequently IFN-γ for efficacy.

**Figure 2.**
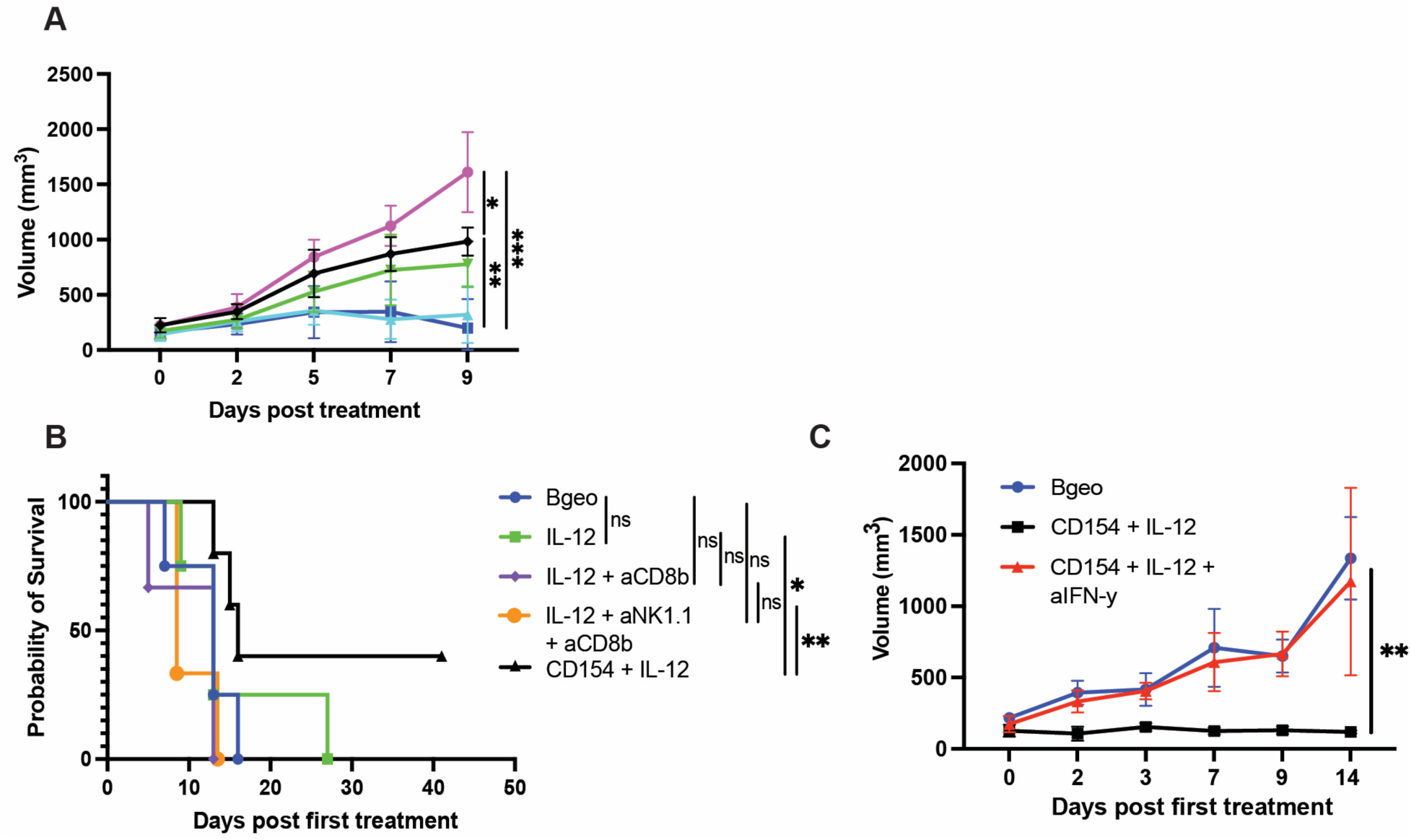
The addition of CD154 to IL-12 electroporation requires CD8 T cells for anti-tumor efficacy. A) Selectively depleting CD8 T cells abrogates the efficacy of CD154 + IL-12 electroporation. B) Similarly, CD8 T cell depletion negates the improved survival with IL-12 only electroporation. C) IFN-γ neutralization eliminates CD154 + IL-12 efficacy. All experiments were repeated at least once with similar results except IFN-γ neutralization. N=4-5. Growth curves were analyzed using two-way ANOVA with p > 0.05 as ns, p < 0.05 as *, p < 0.01 as **, and p < 0.001 as ***.

### CD154 + IL-12 induces more tumoral CD8 T cells that express less PD-1

Because both IL-12 and CD154 + IL-12 rely on CD8 T cells for efficacy, but the combination treatment is superior, we hypothesized that there may be phenotypic difference in the CD8 T cell populations between the two treatments that contributes to tumor clearance. We utilized flow cytometry to evaluate the CD8 T cell population eight days after one treatment for all groups. We observed that CD154 + IL-12 treatment produced more CD8 T cells in the tumor than any other group while IL-12 generated more CD8 T cells than PBS or β-geo (**Figure 3A, B**), however they did not express an increased activation profile or heightened expression of cytolytic proteins (**Supplementary Figure 4A–E)**. While phenotypically the T cells were generally similar across the groups, we observed that there were more CD44hi PD-1+ CD8 T cells in the IL-12 and CD154 + IL-12 treated groups, indicating increased antigen experience. Of those CD44hi PD-1+ antigen-experienced cells, CD154 + IL-12-treated cells expressed far less PD-1 on their surface than similar cells from any other group, alluding to a possibly less exhausted T cell state (**Figure 3C, D)**. Taking this into account, we investigated the effect of PD-1 checkpoint blockade with CD154 + IL-12 therapy (**Supplementary Figure 5A)**. The addition of PD-1 blockade to CD154 + IL-12 did not significantly increase overall survival (**Supplementary Figure 5B, C)**. As compared to all other groups, CD154 + IL-12 treatment results in a larger, less-exhausted CD8 T cell pool.

**Figure 3.**
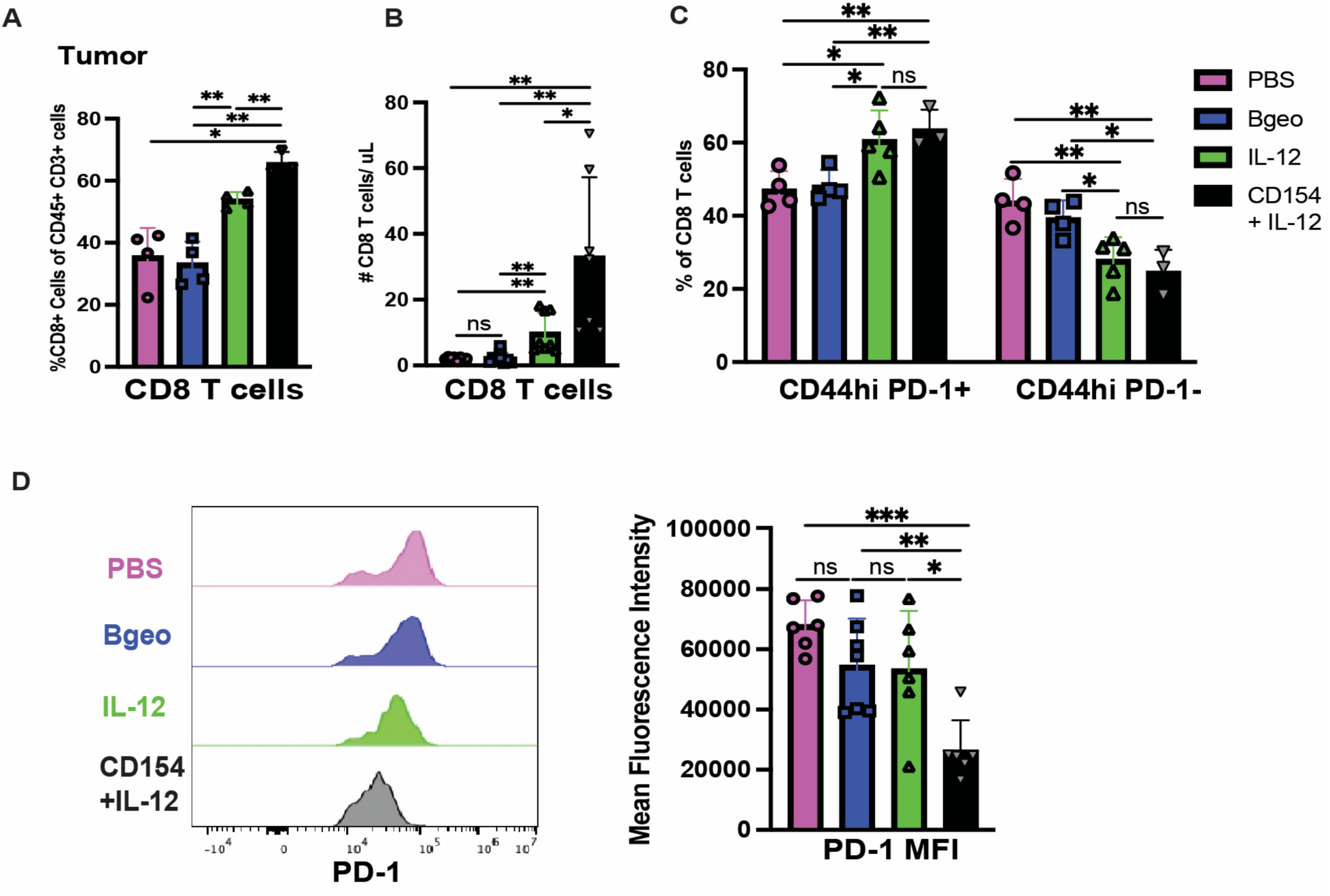
CD154 and IL-12 electroporation generates a greater CD8+ T cell response. A) Tumors treated with CD154+IL-12 electroporation had higher proportion of CD8 T cells among CD45+ cells and B) higher number of CD8 T cells per microliter of sample in the tumor than all other groups. C) A higher frequency of IL-12 and CD154 + IL-12 treated CD8 T cells were CD44hi and PD-1+. D) However, CD154 + IL-12 CD8 T cells expressed less PD-1. Samples were harvested eight days after first treatment and processed for flow cytometry. All experiments were repeated at least once with similar results. B, D represent concatenated results from multiple experiments. Student’s t-test was used to compare the statistical difference between two groups with p > 0.05 as ns, p < 0.05 as *, p < 0.01 as **, and p < 0.001 as ***.

### Dendritic cells treated with CD154 + IL-12 prime more antigen-specific T cells

Since CD154 + IL-12 creates a larger CD8 T cell compartment in the tumor (**Figure 3A, B**), we hypothesized that CD154 + IL-12 generates more antigen-specific T cells. First, we utilized a mismatched tumor model utilizing B16F10 and MC38 to test for systemic T cell antigen specificity. All mice had treated B16F10 tumors with either B16F10 (matched) or MC38 (mismatched) as the non-treated tumors. We observed a significant difference in the matched model on the non-treated tumor, while the mismatched model did not exhibit any difference in tumor growth on the non-treated flank, indicating the generation of an antigen-specific response (**Figure 4 A-C)**. To further investigate CD154 + IL-12 ability to expand antigen-specific CD8 T cells, we utilized Pmel mice, a trackable transgenic CD8 T cell model with a T cell receptor specific for gp100, a melanoma antigen. [33] Mice were given 10,000 naive Pmel CD8 T cells a day before tumor cell inoculation and were treated with plasmid electroporation (**Figure 4D)**. CD154 + IL-12 treated tumors had a significant increase in Pmel cells in tumors, while the tumor-draining lymph node had elevated numbers of Pmel cells in IL-12 and CD154 + IL-12 mice (**Figure 4E-F)**. Due to this increase in antigen-specific T cells, we investigated dendritic cell populations in the tumor draining lymph node to determine if there was a potential difference in ability to prime T cells. CD154 + IL-12 increased the number of dendritic cells in the tumor-draining lymph node but did not affect the proportion of conventional type 1 dendritic cells (defined as XCR1+ CD11c+ MHCII+) which cross prime CD8 T cells among all DCs (**Figure 4G, H).** However, cDC1s in tumor-draining lymph nodes of CD154 + IL-12 treated mice exhibited a heightened CD86 and increased MHCII expression, indicating an enhanced priming capacity (**Figure I, J)**. To validate the importance of cDC1s and their ability to prime the CD8 T cell response in CD154 + IL-12 therapy, we evaluated CD154 + IL-12 in the BatF3-/- mouse model, which lacks cDC1s. In the Batf3-/- mice, CD154 + IL-12 lost its antitumor efficacy, demonstrating the need for cDC1s for tumor response (**Figure 4 K, L).** The data supports the conclusion that adding CD154 to IL-12 electroporation improves cDC1 cross priming of CD8 T cells, leading to an increase in antigen-specific effector CD8 T cells in the tumor.

**Figure 4.**
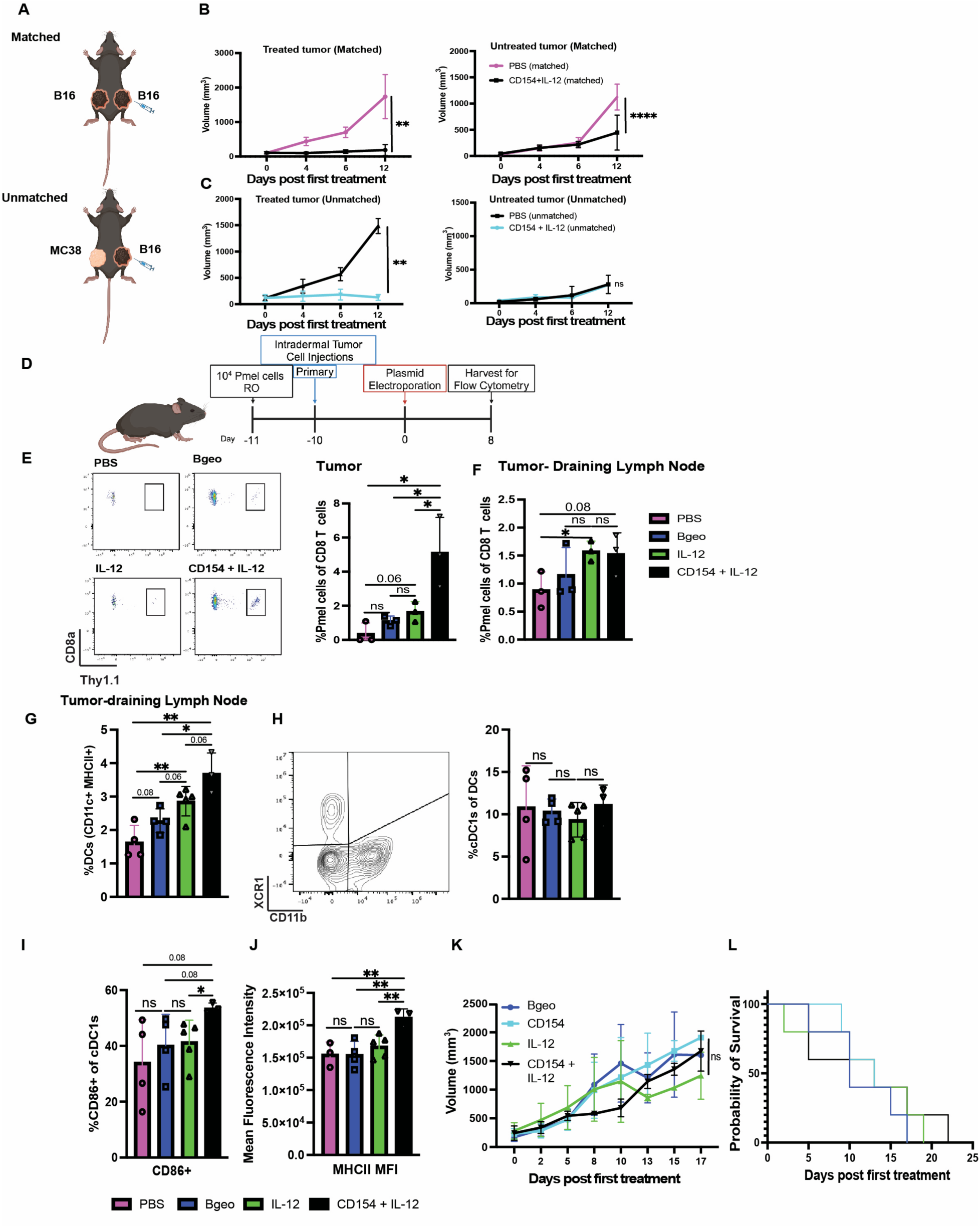
CD154 and IL-12 electroporation increases the number of tumor antigen-specific T cells and induces better dendritic cell priming. A) Schematic for mismatched tumor model. All mice had B16F10 on their treated flank with the non-treated flank being either B16F10 (matched) or MC38 (mismatched). B,C) CD154 + IL-12 only affected matched untreated tumor, while the mismatched untreated tumor showed was not inhibited by treatment. D) One day before tumor inoculation, mice were given Pmel cells retro-orbitally, then received plasmid electroporation. On Day 8, tissues were collected and processed for flow cytometry. E) CD154 + IL-12 induced a higher frequency of antigen-specific (Pmel) T cells in the tumor, F) tumor-draining lymph node had a slight increase in Pmel cells in IL-12 and CD154 + Il-12 compared to control. G) Dendritic cells were increased in the tumor-draining lymph node of CD154 + IL-12 treated mice. H) While the frequency of cDC1s did not change, CD154 + IL-12 treated mice induced a I) higher frequency of CD86+ cDC1s with J) an increased expression of MHCII. K) In Batf3-/- mice, lacking cDC1s, CD154 + IL-12 neither inhibits tumor growth nor L) improves survival. N=4-5. All experiments were repeated at least once with similar results. Student’s t-test was used to compare the statistical difference between two groups. Growth curves were analyzed using two-way ANOVA. Survival curves were compared using log-rank (Mantel-Cox) test, with p > 0.05 as ns, p < 0.05 as *, p < 0.01 as **, and p < 0.001 as ***.

### CD154 + IL-12 reduces T regulatory cells in treated tumors

We determined the CD4 T cell numbers in tumors and observed a 2-fold decrease in CD4 T cells among total T cells in the CD154 + IL-12 mice (**Figure 5A**). This CD4 T cells reduction was most pronounced in T regs. Gating on either FoxP3 and CD25 or FoxP3 expression alone, we observe a decrease in the frequency of FoxP3+ CD25+ T regulatory cells (Tregs) and FoxP3+ cells in both IL-12 and IL-12 + CD154 groups. While reduced in IL-12 treated mice, the number of Tregs in CD154 + IL-12 was even more reduced (**Figure 5 B-D**). Combined with CD8 T cell numbers, the CD8: Treg ratio was increased many fold by CD154 + IL-12 therapy (**Figure 3B, 5E)**. Furthermore, we found that the depletion of CD4 T cells, which depletes T regs as well as other CD4 T cells, when combined with IL-12 electroporation, recapitulated the tumor clearance efficacy of CD154 + IL-12 treatment, indicating that the addition of CD154 improves tumor clearance predominantly through the reduction of Tregs (**Figure 5F)**.

**Figure 5.**
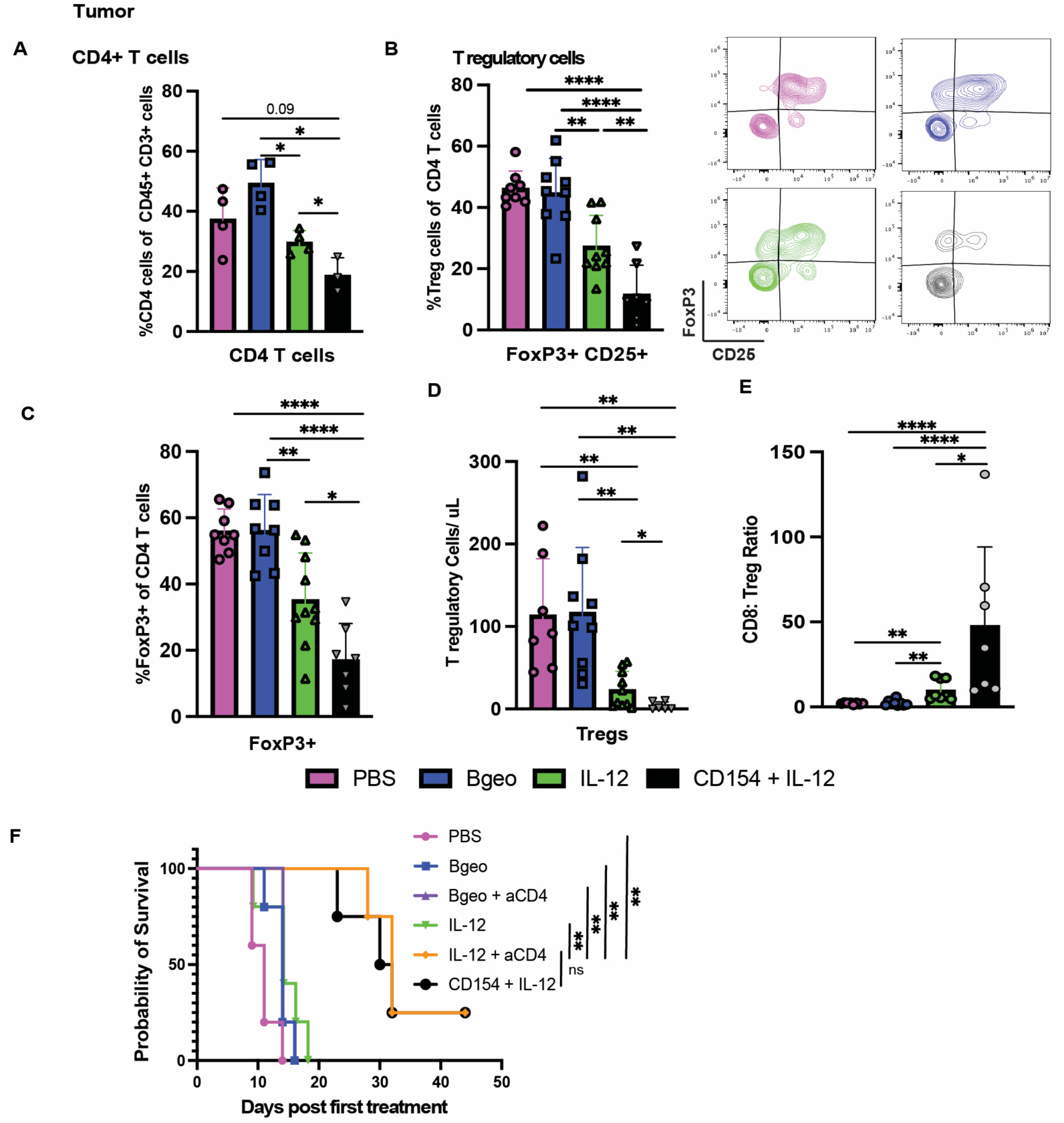
CD154 + IL-12 dramatically decreases intratumoral T regulatory cells. A) Eight days after a single treatment, CD4 T cell frequency decreases significantly in the treated tumor in CD154 + IL-12 treated mice. B) FoxP3+ CD25+ T regulatory cells are a lower % of CD4+ T cells in the tumors of CD154 + IL-12 treated mice than in all other groups. C) Similar results were seen when gating via FoxP3 expression alone on CD4 T cells. D) The decrease in T-reg frequency corresponded with a drastic decrease in cell number of Tregs in tumors treated with CD154 + IL-12. E) CD8 T cell: Treg ratio is increased many fold in CD154+ IL-12 treated tumors. F) For anti-CD4 antibody depletion, mice received two plasmid electroporation treatments a week apart as in all efficacy studies and were given 200ug of anti-CD4 antibody intraperitoneally every three days starting on the day prior to treatment. IL-12 electroporation in conjunction with CD4 T cell antibody depletion recapitulates CD154 +IL-12 efficacy. All experiments were repeated at least once with similar results. B-E represent concatenated results from multiple experiments. Student’s t-test was used to compare the statistical difference between two groups. Survival curves were compared using log-rank (Mantel-Cox) test, with p > 0.05 as ns, p < 0.05 as *, p < 0.01 as **, and p < 0.001 as ***.

## Discussion

The combination of CD154 and IL-12 amplifies changes mediated by IL-12 alone. While IL-12 has widespread effects on various immune effector cells, previous studies of IL-12 electroporation, (reproduced here), identified CD8 T cells as the primary responder to increased IL-12 expression in the TME. [12,34] Similarly, in treated tumors, consistent with other IL-12 electroporation studies, NK depletion offered a slight effect on the response against melanoma tumors, whereas depletion of CD8+ T cells eliminated IL-12-driven [16] and CD154 + IL-12- driven anti-tumor responses, as we show. [15] Initial studies of IL-12 electroporation observed an increase in CD8 T cell infiltration into the TME and the generation and expansion of antigen-specific CD8+ T cells. [15] Follow-up studies involved the introduction of T cell stimulating cytokines to further enhance the CD8 T cell population. [11] Studies utilizing CXCL9, IL-2, TNF-a, anti-CD3, and IL-27 in combination with IL-12 electroporation displayed increased tumor clearance that correlated with increased T cell activity in the TME. [16,22,23,35,36] We acknowledge that initial and subsequent IL-12 studies have observed tumor clearance in B16F10 with IL-12 electroporation only. In our hands, we do not observe clearance with IL-12 alone and likely different experimental conditions explain that. This may be due to a difference in the IL-12 construct, frequency of treatment, electroporation efficiency or residual LPS levels. We utilized a dual prong probe attachment for our electroporation unit (BTX Harvard Gemini X2 electroporator) and expect improved electroporation efficiency and therefore efficacy with a multi-pronged prong probe as used in other studies. [12,16,22,35]

Activation of CD40 on DCs aids in overcoming peripheral tolerance to generate robust anti-tumor response via cross-presentation of antigen to CD8 T cells. [37] However, the timing of CD40 activation on DCs is important relative to antigen uptake. Immature DCs, that are non-activated, exhibit superior antigen uptake and loading, whereas CD40-activated DCs have reduced antigen-loading capacity. Pre-activation with CD40 antibodies prior to antigen availability results in a reduced or even abolished T cell response due to inefficient priming. [38,39] Therefore, most studies suggest that vaccination before CD40 activation or concomitant antigen delivery is optimal. Our study circumvents this timing challenge through the delivery of electrical charge that generates tumor cell death and subsequent tumor antigen release, allowing for tumor antigen uptake by DCs prior to plasmid-driven CD154 expression. Additionally, CD154 expression allows pre-existing immature, tumor antigen-loaded DCs in the tumor to mature and migrate to the draining lymph node, resulting in improved priming and enhanced CD8 T cell response.

The addition of CD154 greatly decreases the Treg population in the TME. Other CD40 agonist studies have also observed a reduction of Tregs. [27,40,41] However, the mechanism remains unclear. In transplant models, CD154 blockade but not CD40 blockade preferentially increased inducible FoxP3+ Tregs in transplanted tissues. The discrepancy in outcome when blocking CD154 versus CD40 indicates that CD154 is potentially binding with another ligand other than CD40. Furthermore, IL-1b blockade in CD40-/- recipients further elevated Treg numbers. [42] Recent studies in alloimmunity have shown that CD11b is an alternative ligand for CD154 through the Mac-1 integrin. [43] Therefore, if the lack of CD154 binding increases Treg induction potentially the inverse is also true and that the over expression of CD154 or increase in CD40 agonism may decrease Tregs, potentially through IL-1b. A future direction would be to investigate the role of CD11b+ cells and IL-1b in CD154 + IL-12 electroporation to determine its potential role in the decrease in tumoral Tregs.

Agonistic CD40 antibodies as a monotherapy in clinical trials have shown a wide range of demonstrated activity. Selicrelumab, the most extensively studied anti-CD40 monoclonal antibody, exhibits a moderate objective response rate in patients with melanoma but had little clinical activity in patients with advanced cancers in other studies. [41,44] Due to the modest clinical response of CD40 monoclonal antibodies, CD40 antibodies are mainly being tested in conjunction with other cancer drugs as a means of increasing response. [41] Another limitation is systemic injection of human agonistic CD40 antibodies have been reported to trigger severe adverse effects and toxicities including hepatotoxicity, cytokine release syndrome (CRS), thrombocytopenia general hyperimmune stimulation and tumor angiogenesis. [29] Therefore, the local CD40 agonism from CD154 electroporation ITIT offers an alternative that limits systemic exposure to CD40, avoiding potential toxicities.

Overall, our results demonstrated that the addition of CD154 to IL-12 electroporation enabled elimination of 30-80% of treated tumors in B16F10 and MC38 models. Clinically, studies involving IL-12 electroporation have shifted away from it as a monotherapy but as a platform for combination immunotherapies, including Immune checkpoint blockade. Similarly, recent publications have harnessed different immunostimulatory cytokines with IL-12 to generate a more robust CD8 T cell response. Considering, the proven clinical safety and efficacy of ITIT IL-12, the potential of adding other immunostimulatory cytokine plasmids to the IL-12 platform should be a priority of cancer immunotherapy research. Concurrently, agonistic CD40 antibody immunotherapy has shown efficacy across multiple tumor types, however, its momentum has severely slowed due to fear of severe adverse effects. We show here that combined intratumoral expression IL-12 and CD154 in tumors nearly eliminated intratumoral T regs, leading to tumor frequent clearance. While reduction in the Treg compartment has been shown in both CD40 agonism studies as well as from IL-12, the extent of T regs elimination by the combination was almost complete. While the concept of targeting T regs in tumors is not new, a deeper understanding of Treg biology and how CD40 ligand depletes Tregs may offer insights into the development of better tumor immunotherapies.

## Methods

### Cell lines

The B16F10 (B16) melanoma cells were provided by coauthor Dr. Mary Jo Turk. MC38 colon adenocarcinoma cell line was purchased from the American Type Culture Collection. B16 cells were cultured in RPMI (Corning) (Corning, NY) supplemented with 10% fetal bovine serum, 2 mM l-glutamine, 0.5% penicillin/streptomycin. MC38 cells were cultured in Dulbecco’s modified MEM (Corning) (Corning, NY) supplemented with 10% fetal bovine serum, 2 mM l-glutamine, 0.5% penicillin/streptomycin. These cell lines were authenticated by morphology, phenotype, and growth, and routinely screened for Mycoplasma, and were maintained at 37°C in a humidified 5% CO2 atmosphere.

### Mice

Female C57BL/6 mice were purchased from the Charles River Laboratories (Wilmington, MA). TLR9 −/− and Batf3−/− mice on C57BL/6 mice background were purchased from the Jackson Laboratories (Bar Harbor, ME) and bred in house, and Rag2 knockout mice were gifts of Professor Yina Huang of Dartmouth Geisel School of Medicine. All mice were age matched (7–12 weeks old) at the beginning of each experiment and kept under specific pathogen-free conditions and housed in the Laboratory Animal Resources. All animal studies were approved by the Institutional Animal Care and Use Committee of Dartmouth College under approved protocol 2137 and followed the ARRIVE guidelines.

### Plasmid purification

Three different plasmids were used in the experiments: IL-12-encoding plasmid was a gift from Prof Shulin Li of University of Texas MD Anderson Cancer Center, CD154-encoding plasmid was purchased from Origene (Rockville, MD) and pβ−geo was from LA Herzenberg, Stanford University, which was used as a control plasmid. Plasmids were amplified in E. coli and purified using Endo Free Plasmid Mega Kits (Qiagen, Hilden, Germany) according to the manufacturer’s protocol and dissolved in endotoxin-free water (Qiagen, Hilden, Germany). The identity of each plasmid was verified by restriction analysis and subsequent agarose gel electrophoresis for the separation of formed bands. The concentration and purity of isolated plasmids was determined using a Nanodrop 2000 spectrophotometer (Thermo Fisher Scientific, Waltham, USA), which measured the 260/230 and 260/280 absorbance ratios.

### Pmel cell adoptive transfer

Naïve CD8+Thy1.1+ pmel cells were isolated from 8–10-week-old pmel T cell receptor transgenic mice. Cells were magnetically sorted by Mouse Naïve CD8 T cell extraction kit (Stemcell Technologies). T cells were transferred retro-orbitally, at 10^4^ cells per mouse, 1 day prior to intradermal tumor inoculation.

### Tumor inoculation

B16F10 (2×10^5^) or MC38 (2×10^5^) tumor cells were intradermally injected in 30 μL Hank’s Balanced Salt Solution (HBSS) on one or both flanks under anesthesia with isoflurane. For two-tumor mouse models, the treated tumor was inoculated on Day −10, the untreated tumor on Day −7 and the treatment was started on Day 0. Tumor growth was monitored every other day. Tumor progression was monitored by measuring tumor volume ((L)^2^ x (W/2)). Mice were euthanized when tumor size reached 1500 mm^3^ or survival reached Day 40. Dermal tumors are optimal for electroporation since the injection and electroporation can be clearly visualized and therefore treatment is more uniform as compared to subcutaneous tumors, which cannot be visualized without opening the skin. Our studies used only female mice because skin tumors are more variable in male mice because their skin is thicker and tougher, and they sometimes damage their own or cage mate tumors, making size assessment less accurate.

### Plasmid electroporation

Intradermal tumors were electroporated with 50 ug of CD154-expressing and/or IL-12-expressing plasmid or control plasmid suspended in sterile water and tumor growth was monitored. Mice were electroporated at 1500v/cm for 6 pulses at 100us durations using BTX Harvard Gemini X2 electroporator with dual-prong 10mm gap electrode. Treatment was given twice one week apart.

### *In vivo* antibody treatment

For in vivo depletion of lymphocytes, 200μg of anti-CD4 antibody (clone GK1.5), (bioreactor supernatants from hybridoma cell lines from Dr. Mary Jo Turk), Anti-CD8β (clone Lyt 3.2, Bio X Cell) and anti-NK1.1 (clone PK136, Bio X Cell) were injected i.p. every third day three times from the first treatment day. For in vivo depletion of IFN-γ, 1 mg of anti-IFN-γ (clone R4−6A2, Bio X Cell) Abs were administrated by i.p. injection at the first treatment day, with follow-up doses of 500μg for five consecutive days. For PD-1 blockade, 200μg of aPD-1 Ab (clone RMPI-14, Bio X Cell) were given intraperitoneally (i.p.) every third day from the first day ITIT.

### Tissue processing

All specimens from a single subject were processed on the day of collection. Tumors were weighed, and then minced with surgical scissors. Minced specimens were dissociated utilizing GentleMACs dissociator (Miltenyi Biotec) and then digested in 5ml RPMI-1640 (Hyclone) containing 1 mg/mL collagenase type IV, at 37 °C for 20 mins, after which an equal volume of cold flow buffer (PBS+2% FBS) was added. Samples were then filtered through a 70 μm cell strainer. The single cell suspension was centrifuged at 1,800 RPM for 5min and resuspended in PBS. Spleens were mashed through a 70 μm cell strainer, spun down and resuspended in RBC Lysis Buffer (Biolegend) at room temperature for 15 mins, after which the lysis was stopped by adding 10-fold volume of PBS, and cells were centrifuged at 1,800 RPM for 5min and resuspended in PBS. Lymph nodes were treated as splenic samples but were not subject to RBC lysis.

### Flow cytometry

Single-cell suspensions of mouse lymph nodes were prepared for flow cytometric analysis. Fc receptors were blocked with anti-mouse CD16/32 (BioLegend) for 15 minutes on ice and surface stained with indicated markers. Flow antibodies used were as follows: NK1.1(PK136), Live Dead fixable blue (Invitrogen), CD11c (N418), CD45(30-F11), Ly6C (HK1.4), CD8a (53-6.7), XCR1 (ZET), CD103 (M290), PD-L1 (10F.9G2), Ly6G (1A8), CD11b (M1/70), CD3(17A2), CD86 (PO3), F4/80 (BM8), MHCII (M5/114.15.2), CD19 (6D5), CD154 (MR1), CD44 (IM7), PD-1 (29F.1A12), Granzyme B (QA16A02), Perforin (S16009A), CD69 (H1.2F3), FoxP3 (150D), Thy1.1 (OX-7) and CD25 (PC61). After 30 min incubation on ice, cells were washed twice with Flow buffer (PBS with 2% FBS) and were fixed with 4% Paraformaldehyde for 20 minutes. For intracellular antibody staining, we used an intracellular FoxP3/ transcription factor staining kit (Tonbo Biosciences) samples were resuspended in fixation/permeabilization buffer incubated for 40 min at room temperature in the dark, and then washed with permeabilization buffer. Samples were resuspended in 100 μl of permeabilization buffer, and fluorescently labeled antibodies against intracellular targets were added to the appropriate samples. Samples were incubated for 40 min at room temperature in the dark and then washed twice with permeabilization buffer. Samples were then fixed in 4% Paraformaldehyde as stated above. We used the published gating strategy to dissect the myeloid and lymphoid compartment changes with CYTEK Aurora Spectral cytometer and FlowJo V.10.10.0.

### Statistical analysis

All results are expressed as means ± SEM (n=3–5) as indicated. Student’s t-test was used to compare the statistical difference between two groups. Growth curves were analyzed using two-way ANOVA. Survival curves were compared using log-rank (Mantel-Cox) test, with p > 0.05 as ns, p < 0.05 as *, p < 0.01 as **, and p < 0.001 as ***. GraphPad Prism V.10.2.2 software was used to calculate significance between the samples. FlowJo V.10.10.0 was used to analyze corresponding data. All experiments were repeated, and representative figures were presented unless otherwise noted. P values≤0.05 were considered significant.

## Declarations

### Contributors

**Gregory Ho,** conceptualization, methodology, investigation, writing – original draft, review and editing. **Alicia Santos**, investigation. **Jennifer Field**, investigation. **Pamela C. Rosato**, writing – review and editing**. Mary Jo Turk**, writing – review and editing. Nicole **F. Steinmetz:** funding acquisition, writing – review and editing. **Steven Fiering**: supervision, project administration, funding acquisition, writing – review and editing.

### Funding

This research was funded by R01 CA224605 to NFS.

### Competing interests

SF and NSF are co-founders of, have equity in, and have a financial interest with Mosaic ImmunoEngineering Inc. NFS serves as manager of Pokometz Scientific LLC, under which she is a paid consultant to Flagship Labs 95 Inc., and Arana Biosciences Inc. The other authors declare no potential conflicts of interest.

### Patient consent for publication

Not applicable.

### Ethics approval

All procedures and animal experiments were reviewed and approved by the Institutional Animal Care and Use Committee (IACUC) at Dartmouth College.

### Data Availability Statement

All data relevant to the study are included in the article or uploaded as online supplemental information. Any further information about resources and reagents should be directly requested to the corresponding authors and will be fulfilled on reasonable request.

## Acknowledgements

This study was supported by Dartmouth Mouse Modeling Shared Resource, Preclinical Imaging and Microscopy Shared Resource and Dartlab Immune Monitoring Shared Resource, which receive support from the Dartmouth Cancer Center, through Funding from NCI P30CA023108.

